# Condensed chromatin behaves like a solid on the mesoscale in vitro and in living cells

**DOI:** 10.1101/2020.05.06.079905

**Authors:** Hilmar Strickfaden, Thomas Tolsma, Ajit Sharma, D. Alan Underhill, Jeffrey C. Hansen, Michael J Hendzel

## Abstract

The association of nuclear DNA with histones to form chromatin is essential to the temporal and spatial control of eukaryotic genomes. In this study, we examined the physical state of chromatin in vitro and in vivo. Our in vitro studies demonstrate that MgCl2-dependent self-association of native chromatin fragments or reconstituted nucleosomal arrays produced supramolecular condensates whose constituents are physically constrained and solid-like. Liquid chromatin condensates could be generated in vitro, but only using non-physiological conditions. By measuring DNA mobility within heterochromatin and euchromatin in living cells, we show that chromatin also exhibits solid-like behavior in vivo. Representative heterochromatin proteins, however, displayed liquid-like behavior and coalesced around a solid chromatin scaffold. Remarkably, both euchromatin and heterochromatin showed solid-like behavior even when transmission electron microscopy revealed limited interactions between chromatin fibers. Our results therefore argue that chromatin is not liquid but exists in a solid-like material state whose properties are tuned by fiber-fiber interactions.

## INTRODUCTION

The structural organization of chromatin in the nucleus remains poorly understood (Bian and Belmont, 2012; Eltsov et al., 2008; Fussner et al., 2011; Joti et al., 2012; Maeshima et al., 2010; Maeshima et al., 2016b; Ou et al., 2017). Long standing hierarchical models of chromatin folding have largely been disproven and new models based on clustering of nucleosomes (Otterstrom et al., 2019; Ricci et al., 2015) and extended 10 nm chromatin fibers (Fussner et al., 2012; Maeshima et al., 2010; Maeshima et al., 2016b) into compact chromatin domains (Hansen et al., 2018) have taken their place. In these new models, interphase and mitotic chromosomes are assembled from 10 nm chromatin without subsequent helical coiling. As a consequence, the classical 30 nm fiber that was widely thought to be essential to chromosomal folding and transcriptional repression (Hansen, 2002), is now thought to be absent from most animal cells and tissues. While the evidence in favor of models lacking the 30 nm fiber is substantial, the mechanisms by which extended 10 nm fibers are packaged into higher order heterochromatin and euchromatin domains have not been established. An important aspect of this question is whether chromatin is mobile and liquid-like physically more constrained within chromosomal environments, because it is critical to understanding how genome function is regulated. Concomitant with the new models of chromatin organization in the nucleus, in vitro studies of chromatin dynamics also have shifted focus. After decades of focusing on 30 nm fibers (Hansen, 2002; Thoma et al., 1979), recent studies have investigated the process by which chromatin condenses into supramolecular aggregates(Gibson et al., 2019; Maeshima et al., 2016b; Maeshima et al., 2020), which we will refer to as chromatin condensates. Folded 30 nm structures are first observed when cations are titrated into a purified chromatin sample (Finch and Klug, 1976; Thoma et al., 1979). At higher cation concentrations that better approximate those found in vivo (Strick et al., 2001), chromatin assembles into condensates. Chromatin condensates have been studied for many years. Historically, chromatin condensates were referred to as precipitants or insoluble aggregates based on the ease at which they pelleted upon centrifugation. Nevertheless, condensate assembly is freely reversible and requires the core histone tail domains (Gordon et al., 2005; Schwarz et al., 1996), suggesting chromatin self-interaction is biologically relevant. The structural features of chromatin condensates formed by defined 12-mer nucleosomal arrays in 4-10 mM MgCl2 have recently been determined (Maeshima et al., 2016b). The condensates had a globular morphology and reached 0.5-1.0 micron in diameter as judged by fluorescence and electron microscopy. Moreover, sedimentation velocity experiments indicated that the condensates sedimented in the 300,000S range, thereby confirming their size in solution (Maeshima et al., 2016b). The individual nucleosomal arrays that made up the condensates were packaged as irregular 10 nm fibers, as is the case in vivo (Fussner et al., 2012). One property that was not determined by Maeshima et al. (2016b), however, was the mobility of packaged nucleosomal arrays within the condensates. This question was recently addressed by Gibson et al. (2019), who studied cation-driven condensate formation in the presence of acetate, dithiothreitol (DTT) and bovine serum albumin (BSA) was best described as a liquid/liquid phase separation process. The authors observed chromatin mobility within condensates, which they attributed to liquid/liquid phase separation, leading them to speculate that this also represents the material state of chromatin in vivo.

Despite the attractiveness of liquid condensate formation as a mechanism to explain chromatin packaging and condensation, the contribution of chromatin to the mechanical integrity of the nucleus and its ability to respond to extranuclear forces applied to the nuclear surface are difficult to reconcile with a liquid state. Two groups have recently studied the contribution of chromatin organization to the mechanical stability of the nucleus. One found that the ability of the nucleus to resist an applied force was reduced approximately 10-fold by mildly digesting linker DNA and approximately 3-fold by treating cells with histone deacetylase inhibitors (Maeshima et al., 2018), indicating that intact chromatin is needed to maintain the structural integrity of the nucleus. Marko and colleagues further demonstrated that cells adapt to mechanical strain through activation of mechanically sensitive ion channels, which stimulates the histone methylation machinery to drive the assembly of more heterochromatin. The increased heterochromatinization of the nucleus resulted in an increased resistance to externally applied stress (Stephens et al., 2019). These results demonstrate that nuclear chromatin is mechanically responsive and can resist significant applied force. This mechanical property is more consistent with a solid or gel state of bulk chromatin during interphase.

In this study, we critically assessed the mobility of defined nucleosomal arrays and native chromatin fragments within chromatin condensates that form under the conditions of (Maeshima et al., 2016b) and (Gibson et al., 2019) together with directly assessing liquid-like properties of chromatin living cells. When the properties of the condensates formed in vitro in MgCl2-containing buffers were examined, we found that the packaged nucleosomal arrays were immobile, as in a solid- or gel-like state. Liquid condensates could be observed, but only in the presence of DTT, BSA, and acetate anions. Under all other conditions examined, the packaged nucleosomal arrays within the condensates were positionally constrained and solid-like. For our studies of chromatin in vivo, we incorporated fluorescent nucleotides into the genomic DNA of living cells, which enabled tracking of the mobility of the chromosomal DNA molecule rather than the associated proteins. We found that DNA motion within heterochromatin and euchromatin domains was constrained, such that it did not mix with surrounding chromatin even after hyperacetylation-induced decondensation. Finally, using constitutive heterochromatin of the pericentromere, we show that such solid-like chromatin nevertheless supports the formation of dynamic liquid protein assemblies. Taken together, our study provides concordant in vitro and in vivo results that indicate condensed chromatin is packaged into a solid- or gel-like state and addresses the ramifications of a globally constrained genome.

## RESULTS

### Condensed chromatin in vitro is packaged into a solid-like state that is sensitive to disruption by DTT, BSA, and acetate

In vitro, chromatin forms condensates under a wide range of salt conditions (Gibson et al., 2019; Hansen, 2002; Maeshima et al., 2016b; Perry and Chalkley, 1982). The chromatin within the condensates is packaged as irregular 10-nm fibers in a manner that resembles chromatin packaging within chromosomes in vivo (Hansen et al., 2018; Maeshima et al., 2016b). In this study we asked whether the chromatin in the condensates is mobile as in a liquid or constrained as in a solid. We first examined the condensates formed by native chromatin fragments derived from micrococcal nuclease digestion, which consist of a heterogeneous population of nucleosomal arrays bound to H1 and a host of other chromatin-associated proteins. Consistent with a large body of published results (Hansen, 2002; Perry and Chalkley, 1982), native chromatin did not sediment appreciably in 100 mM NaCl but formed pelletable condensates in the presence of added MgCl2 (Figure 1A). Fluorescence microscopy in 100 mM NaCl/2 mM MgCl2 revealed the presence of isolated chromatin condensates that were globular and 0.46 – 0.63 μm in diameter (Figure 1B). Upon contact, the individual condensates formed complex 3D structures rather than fusing into larger spheres (Figure 1B, Movie 1). When the condensates were subjected to FRAP, we observed a lack of mixing of the bleached and unbleached chromatin even after overnight incubation (Figure 1C). The findings that the chromatin condensates formed discrete pellets upon centrifugation (Figure 1A), did not fuse upon contact (Figure 1B), and did not mix after photobleaching (Figure 1C) all indicate that the native chromatin within the condensates was packaged in a constrained solid-like state.

**Figure 1.**
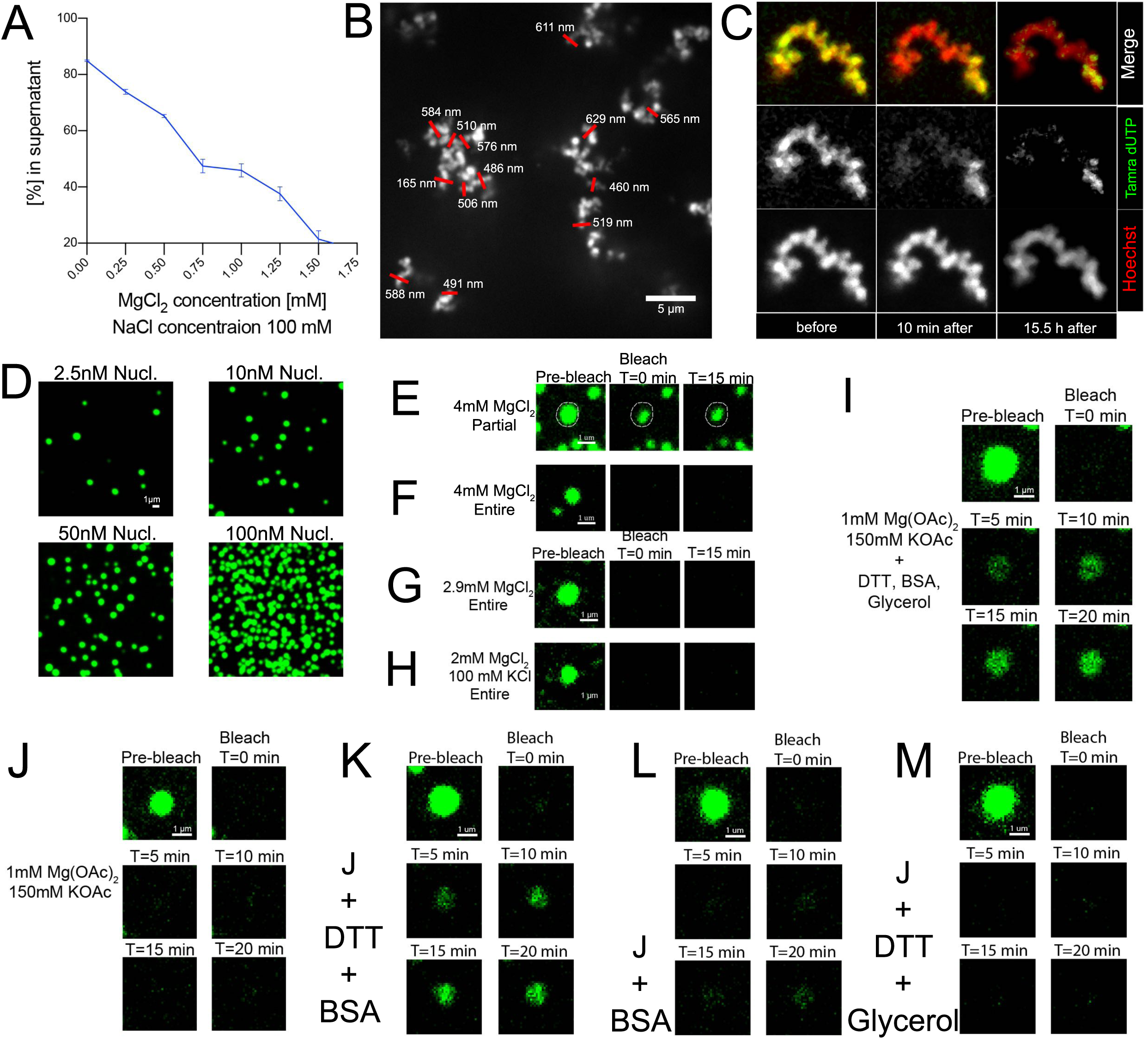
Solid-like behavior of chromatin condensates in vitro. **A:** The formation of pelletable condensates by micrococcal nuclease-digested native chromatin of CH310T1/2 cells in the presence of 100 mM NaCl and increasing concentrations of magnesium was measured by recording the absorbance at 260 nm in solution following centrifugation to sediment the condensates. **B:** Native aggregated chromatin condensates formed in 5 mM MgCl2 were stained with Hoechst and visualized by fluorescence deconvolution microscopy. Full-width half maximum (FWHM) measurements of condensate diameters are shown. **C:** FRAP of native chromatin condensates stained with Hoechst and TAMRA-dUTP immediately after photobleaching, 30 minutes after photobleaching, and 24h after photobleaching. **D:** Condensates formed in 4 mM MgCl2 by Alexa488-labeled 601-207×12 nucleosomal arrays were visualized by fluorescence microscopy. The total nucleosome concentration in the sample is indicated. **E:** Partial FRAP of a 601-207×12 condensate formed in 4 mM MgCl2. **F:** FRAP of an entire 601-207×12 condensate formed in 4 mM MgCl2. **G:** FRAP of an entire 601-207×12 condensate formed in 2.9 mM MgCl2. Approximately 15% of the nucleosomal arrays were unassociated in 2.9 mM MgCl2 compared to 2% in 4 mM MgCl2. **H:** FRAP of an entire 601-207×12 condensate formed in 2 mM MgCl2/100 mM KCl. **I:** FRAP of an entire 601-207×12 condensate formed in 150 mM KOAc/1 mM Mg[OAc]2 plus 5 mM DTT, 0.1 mg/ml BSA, and 5% glycerol. **J:** FRAP of an entire 601-207×12 condensate formed in 150 mM KOAc/1 mM Mg[OAc]2 only. **K:** FRAP of an entire 601-207×12 condensate formed in 150 mM KOAc/1 mM Mg[OAc]2 plus 5 mM DTT and 0.1 mg/ml BSA. **L:** FRAP of an entire 601-207×12 condensate formed in 150 mM KOAc/1 mM Mg[OAc]2 plus 0.1 mg/ml BSA. **M:** FRAP of an entire 601-207×12 condensate formed in 150 mM KOAc/1 mM Mg[OAc]2 plus 5 mM DTT and 5% glycerol.

The solid-like properties of native chromatin condensates were unexpected based on recent descriptions of liquid chromatin condensates formed in vitro by of arrays of positioned nucleosomes (Gibson et al., 2019). It is possible that the presence of H1 or other proteins in our native chromatin was responsible for this discrepancy. We therefore carefully reexamined salt-dependent condensate formation by model nucleosomal arrays. For these experiments, we utilized fluorescently labeled 12-mer 601 nucleosomal arrays with a 207 bp nucleosome repeat length. We first examined the condensates that formed in 4 mM MgCl2, standard conditions used previously for structural analyses (Maeshima et al., 2016b). Greater than 95% of the nucleosomal arrays assemble into pelletable condensates under these conditions (Maeshima et al., 2016b). Consistent with prior results, the condensates under these conditions were globular and ranged from 0.5-1.0 μm (Figure 1D). The condensates did not increase in size with increasing concentration, and at high concentrations individual condensates formed networks of contacting 3D structures rather than fusing into larger globules (Figure 1D), behavior that is inconsistent with a liquid state. We next determined the mobility of the packaged nucleosomal arrays within the condensates using FRAP. When a single condensate formed in 4 mM MgCl2 was partially bleached, there was no mixing of the unbleached arrays and the bleached arrays over a 15 min period (Figure 1E). No fluorescence recovery was observed when an entire condensate was bleached, regardless of whether the free array concentration in solution was very low (in 4 mM MgCl2) or significantly higher (in 2.9 mM MgCl2) (Figure 1F and G), indicating that there was no exchange between the packaged nucleosomal arrays within the condensates and the free nucleosomal arrays in solution. Finally, no recovery was observed for condensates formed in 100 mM KCl/2 mM MgCl2 (Figure1H), a similar condition used to characterize native chromatin condensates (Figure 1B, C). Collectively, the data in Figure 1 demonstrate that formation of chromatin condensates with solid-like properties in the presence of physiological concentrations of MgCl2 and MgCl2/NaCl mixtures is an intrinsic property of arrays of nucleosomes.

The observation of solid chromatin condensates differs from the results of Gibson et al. (2019), prompting us to assess what promoted liquid chromatin under their conditions. We first characterized the condensates that form in 150 mM KOAc/1 mM Mg[OAc]2 plus 5 mM DTT, 0.1 mg/ml BSA, and 5% glycerol, the conditions employed in their study (Gibson et al., 2019). Strikingly, when FRAP was performed on whole condensates formed under these conditions, we observed a linear increase in the level of fluorescence recovery over a 20 min period (Figure 1I; SFig. 1A, Movie 2). These results indicate that the individual nucleosomal arrays within the condensates could diffuse within the condensate as well as exchange with free nucleosomal arrays in solution, indicative of liquid condensates. In view of these findings, we systemically determined which component(s) of the buffers used by (Gibson et al., 2019) were responsible for liquid-like behavior. In 150 mM KOAc/1 mM Mg[OAc]2 buffers without added glycerol, DTT and BSA there was no recovery after photobleaching of entire globules (Figure 1J, Movie 3). Thus, liquid behavior of the condensates required one or more of the additives. In the presence of DTT and BSA, the chromatin remained in a liquid state, indicating that glycerol was not necessary for liquid behavior (Figure1K, Movie 4). When only BSA was present, a very small but measurable amount of recovery was observed after 20 min (Figure 1L, Movie 5). The BSA effect was greatly accentuated in the presence of DTT (Figure 1K, Movie 4). Moreover, when BSA was removed, the condensates did not recover after photobleaching (Figure 1M, Movie 6). We conclude from these results that BSA is absolutely required for liquid-like behavior and that the action of BSA is enhanced by the addition of DTT. This in turn implies that maximum liquidity of the condensates results from the presence of reduced BSA in the buffer. In this regard, reduced BSA is a more potent emulsifying agent than BSA alone (Young Lee and Hirose, 2014) However, to our surprise when DTT and BSA were added to solutions containing 4 mM MgCl2, the condensates did not recover after photobleaching (SFig1B). Thus, liquid chromatin condensates only were observed under a highly specific set of conditions, requiring a combination of acetate anions, DTT, and BSA. Under all other solution conditions tested, the packaged nucleosomal arrays within the chromatin condensates were constrained and solid-like.

### Replication labeling reveals solid-like behavior of euchromatin and heterochromatin in living cells while heterochromatin-associated proteins retain liquid-like behavior

To ascertain whether condensed chromatin in the nucleus is a liquid, the extent of mixing within heterochromatin and euchromatin regions of mouse C3H10T1/2 cells was determined. We first examined the properties of pericentric heterochromatin, which previously has been shown to be associated with membraneless compartments enriched in heterochromatin-associated proteins in Drosophila and murine embryonic fibroblasts termed chromocenters (Strom et al., 2017). If the chromatin within the chromocenters is liquid, photobleaching of labelled DNA would show evidence of diffusion of the label within the chromocenter. To test this, we pulse-labelled cells with a fluorescent nucleotide, TAMRA-dUTP, which was incorporated into the DNA (Schermelleh et al., 2001). When cells are pulse-labelled in S-phase, they give distinctive labeling patterns related to temporal control of genome regulation. In early S-phase, euchromatin replicates and the labelled chromatin is seen as hundreds of individual foci scattered throughout the nucleoplasm (Ferreira et al., 1997). The individual foci are believed to correspond to single topologically associated domains (TADs). Following the replication of euchromatin, the perinucleolar, perinuclear, and pericentric heterochromatin replicate. This enabled us to independently study cells that had incorporated label into heterochromatin or euchromatin. We imaged the cells 24-48 hours after labelling. We observed that the individual chromocenters contained both labeled and unlabeled regions of chromatin, indicating that there was no mixing of the chromatin within the chromocenters over long periods of time (Figure 2A). This is clearly distinct from the behavior of liquid condensates in vitro, where diffusion results in the mixing of labelled and unlabeled macromolecules within the condensate (Hyman et al., 2014; Nakashima et al., 2019). To independently assess the physical state of the chromatin within mouse chromocenters, we performed fluorescence recovery after photobeaching (FRAP) experiments. This is also illustrated in Figure 2A and Movie 7, which show an example where a line is photobleached to bisect several chromocenters and then the labelled DNA is followed over time. Consistent with the lack of mixing of replication-labeled DNA, we found little to no recovery, reflecting limited movement of the chromatin between photobleached and unbleached regions within each chromocenter.

**Figure 2.**
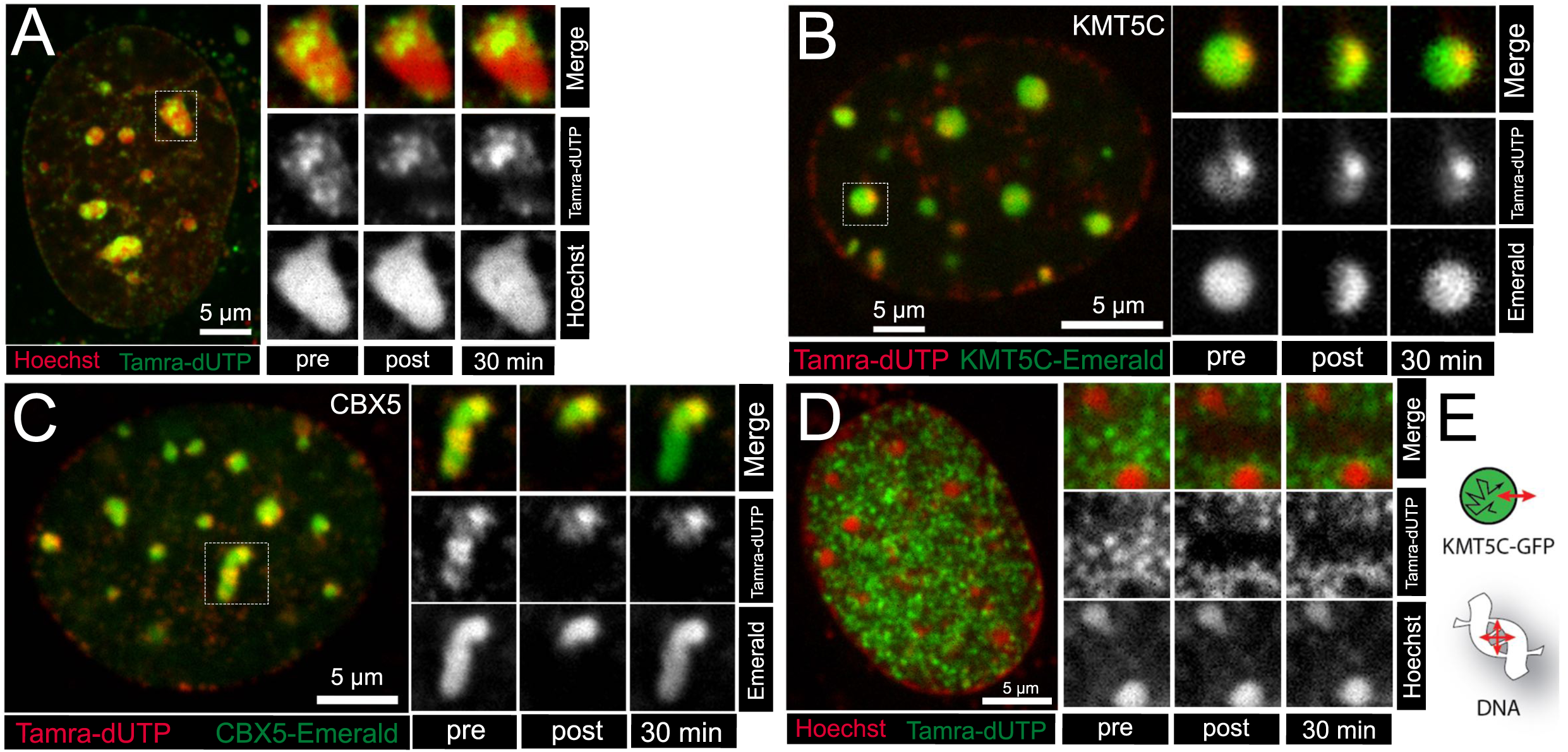
Solid-like behavior of heterochromatic and euchromatic chromatin in living cells. **A**: A CH310T1/2 nucleus stained with Hoechst (red) for DNA and showing chromatin domains labelled with TAMRA-dUTP (green). Insets show the chromocenter (outlined by dashes on the left) before photobleaching, immediately after, and 30 minutes after photobleaching are shown on the right. **B:** Living CH310T1/2 nucleus replication-labelled with TAMRA-dUTP (red) and transfected with KMT5C-Emerald (green). Insets show the chromocenter on the left outlined by dashes before photobleaching, immediately after and 30 minutes after photobleaching (separated by channels and merged). **C:** Living CH310T1/2 nucleus replication-labelled with TAMRA-dUTP (red) and transfected with CBX5-Emerald HP1-alpha (green). Insets show the chromocenter on the left outlined with dashes before photobleaching, immediately after and 30 minutes after photobleaching (separated by channels and merged). **D:** Living CH310T1/2 nucleus stained with Hoechst (red) for DNA and showing chromatin domains labelled with TAMRA-dUTP (green). Insets show an area full of replication foci of early replicating (euchromatin) from the area on the left outlined by dashes before photobleaching, immediately after FRAP and 20 minutes after photobleaching (separated by channels and merged). **E**: Schematic illustrating the mobility of DNA and KMT5C-GFP in a compartment that has undergone liquid-liquid phase separation.

As a control to test whether we could detect phase separated liquid structures in cells where the DNA has incorporated fluorescent nucleotides, we examined two chromocenter-associated proteins that have been reported to be in a liquid state within the chromocenter. KMT5C, a histone H4 lysine 20 methyltransferase, shows liquid behavior within chromocenters but does not exchange with the nucleoplasm (Strickfaden et al., 2019). In contrast, CBX5, which can phase separate at high concentrations in vitro (Larson et al., 2017; Strom et al., 2017), was found to move rapidly into and out of mouse chromocenters (Erdel et al., 2020; Strom et al., 2017). We therefore performed two-channel FRAP experiments on TAMRA-dUTP labelled cells transfected with fluorescent protein tags on KMT5C or CBX5 (Figure 2B, C). We find that KMT5C retains its liquid-like behavior within the chromocenter as evidenced by the very slow but obvious recovery of KMT5C within the partially photobleached chromocenter (Figure 2B, Movie 8, SFig 2A), and CBX5 exchanges rapidly with the nucleoplasm (Figure 2C, Movie 9, SFig 2B), unlike the underlying chromatin which remains positionally stable and does not mix (Figure 2A). Thus, these two heterochromatin proteins, whose diffusion properties support the existence of a liquid unmixed heterochromatin compartment, retain their previously reported kinetic properties in cells that have incorporated TAMRA-dUTP into the DNA. This indicates that the failure to observe liquidity in the labelled DNA is unlikely to reflect a change in the properties of heterochromatin resulting from the incorporation of the modified nucleotide.

We also examined the physical state of euchromatic domains labelled during S-phase. Cells labelled in early S-phase incorporate label into distinct small foci that are thought to correspond to individual topologically associated domains (TADs)(Xiang et al., 2018). We photobleached a region containing several individual foci and followed their fluorescence over time. As shown in Figure 2D and Movie 10, the fluorescence remains depleted over the time course of the experiment. This indicates that chromatin does not mix between individual euchromatic domains. Altogether, the results shown in Figure 2 indicate that both heterochromatin and euchromatin domains in the nucleus exist in a solid-like state that precludes mixing with the surrounding nuclear environment, while protein components that associate with pericentric heterochromatin retain liquid-like properties within the solid domain.

### Acetylation decondenses chromatin but does not induce liquid behavior in living cells

Acetylation weakens the interactions of the core histone tails with DNA (Hong et al., 1993) and was recently reported to promote mixing of microinjected nucleosome arrays with nuclear chromatin (Gibson et al., 2019). Consequently, we next asked whether acetylation could transform solid-like chromatin into liquid chromatin in vivo. To address this question, we treated cells with the histone deacetylase inhibitor Trichostatin A (TSA) to induce hyperacetylation of the histone tails. We first assessed the impact of TSA treatment on the structure of chromatin by direct visualization. Electron spectroscopic imaging (ESI) is a type of transmission electron microscopy that selectively enhances the contrast of phosphorus-rich structures such as chromatin (Hendzel et al., 1999). Using ESI, we determined the distribution of sizes of chromatin-dense structures in 50 nm thick cross sections of interphase nuclei. The images show phosphorus maps (grayscale), where the chromatin is visible, while the color maps show combined nitrogen (red) and phosphorus maps (green), resulting in yellow chromatin and red protein-rich structures. For example, the nitrogen highlights nonchromatin structures such as the nuclear pores (see dashed box in Figure 2B). To provide a sense of scale, circles of 10 nm, 30 nm, and 100 nm diameter are shown. Under control conditions, there are large amounts of dense chromatin structures beyond the heterochromatin found in chromocenters and in association with the lamina. Many of these fibers are ≥100 nm diameter, irregular in shape within the 50 nm thick section, and scattered throughout the nucleoplasm (Figure 3A, - TSA). At higher magnification (Figure 3B), these larger chromatin dense structures have a fibrillar substructure where short sections of fibers of approximately 10 nm in diameter (see ovals in Figure 3B), and consistently less than 30 nm in diameter, can be observed to be packaged together to form the approximately 100 nm diameter chromatin-dense structures (Figure 3B, -TSA). In contrast, apart from chromocenters, the thin sections from cells treated with TSA reveal no obvious chromatin-dense structures throughout the nucleoplasm (Figure 3A, B +TSA). Chromocenters persist under these conditions (Figure 3B chromocenters +TSA) and there may be some retention of higher density chromatin associated with the nuclear lamina (Figure 3A +TSA, 3B Periphery +TSA), but, in general, the chromatin signal is largely punctate (Figure 3A,B), reflecting the dispersal of the dense ∼100 nm chromatin structures found throughout the nucleoplasm. These results are consistent with much of the interphase chromatin, in addition to that found in the chromocenters, is organized into condensed chromatin. It is that widely distributed condensed chromatin found outside chromocenters that is readily decondensed and dispersed by acetylation.

**Figure 3.**
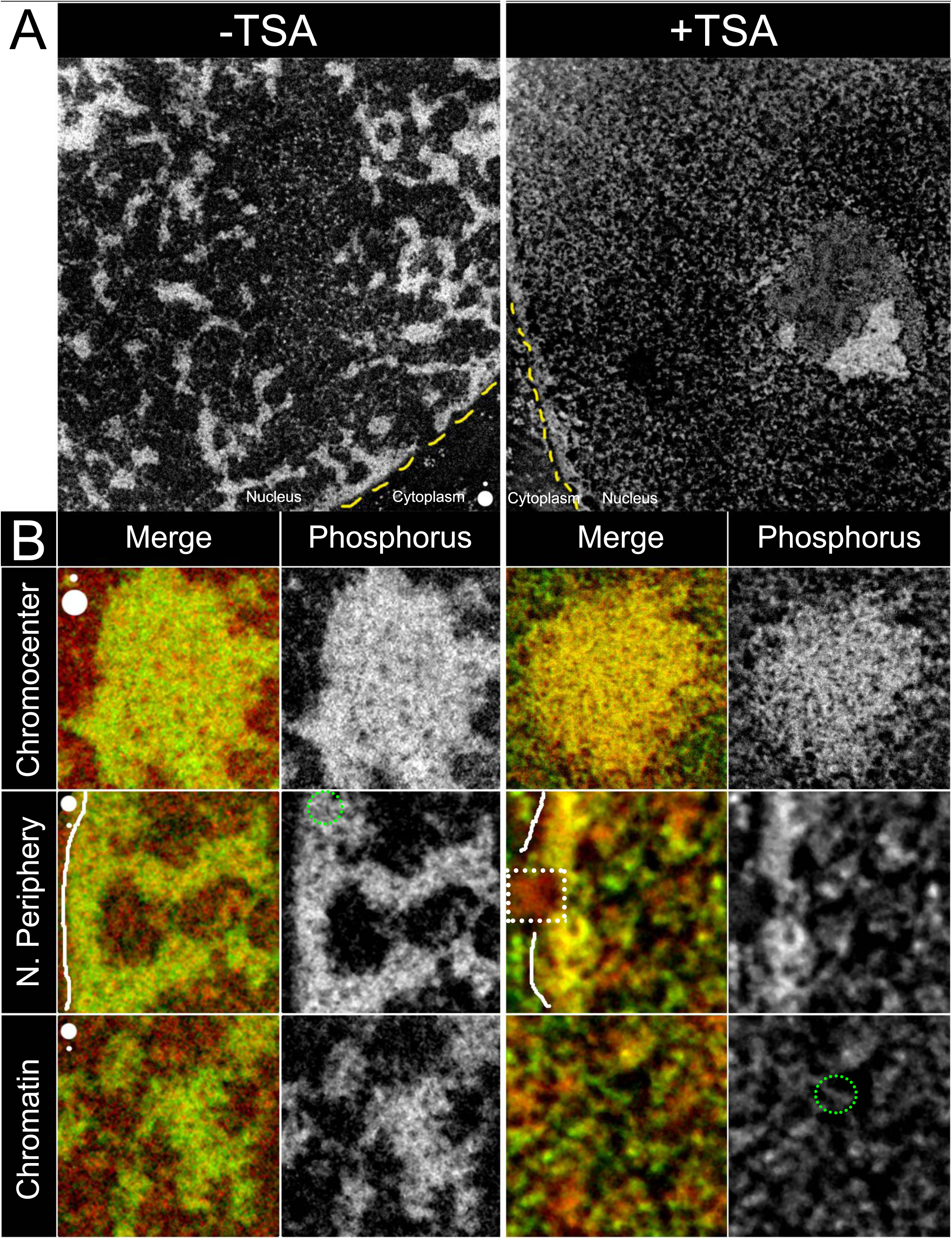
Histone deacetylase inhibition results in global chromatin decondensation. **A**: Transmission electron microscopy images collected by electron spectroscopic imaging are shown. A phosphorous elemental maps of 50 nm ultrathin sections showing a nucleus of a C3H/10T1/2 cell (left) and a nucleus of a cell treated with 100 nM TSA for 24h (right). The circles on the bottom right of the -TSA image are 30 nm and 100 nm in diameter. A portion of the cytoplasm is seen in both images (labelled cytoplasm), where ribosomes can be seen in the phosphorus maps. The dashed yellow lines indicate the boundary between the cytoplasm and the nucleus. **B:** ESI micrographs illustrating chromocenters, peripheral chromatin (N. Periphery) and nucleoplasmic chromatin (Chromatin) in absence (left) and presence (right) of a 24h treatment with 100 nM TSA. The solid line in the N-periphery color images indicate the position of the boundary between the nucleoplasm and cytoplasm. The position of a nuclear pore is highlighted by the dashed box in. For the chromocenter images, the circles represent diameters of 100 nm and 30 nm. In the bottom two rows, the circles represent diameters of 30 nm and 10 nm. The green dashed circles highlight regions where the individual chromatin fibers of approximately 10 nm diameter can be seen.

Having established that, in the presence of TSA, there is minimal dense chromatin formation within the nucleoplasm outside of chromocenters, we assessed the properties of the hyperacetylated chromatin using time lapse microscopy and FRAP. After TSA treatment, the chromocenters persisted (Figure 4A), as expected from the transmission electron microscopy data. Interestingly, despite the observed disassembly of the condensed chromatin found throughout the nucleoplasm, cells where euchromatin incorporated the label during the pulse, the labelled chromatin was still found to persist as numerous independently visible foci (Figure 4B). When these foci were subjected to FRAP, no recovery of the bleached regions occurred over a 30 min period (Figure 4A, Movie 11, 4B, Movie 12). The individual foci, which represent regions of chromatin that have incorporated label at the time of the pulse and are thought to reflect individual TADs (Xiang et al., 2018), are able to move in x, y, and z directions to a limited extent (see Movie 12, see also Movie 10). The individual foci that were fully bleached did not recover significant amounts of fluorescence (Figure 4B), indicating that chromatin does not readily mix between unbleached foci and the foci that were partially bleached. Moreover, while these individual foci show some motion, the failure of the photobleached region to be repopulated by non-photobleached foci surrounding the photobleached region highlights how restricted the motion of even this dispersed chromatin undergoes in living cells. These data are inconsistent with acetylation driving chromatin into a liquid state within cells as observed by Gibson et al., (2019), where the acetylated chromatin readily mixed and dispersed within the nucleoplasm. Rather, our results indicate that there are additional constraints on endogenous chromatin that may not apply to the short microinjected arrays examined by Gibson et al. (2019).

**Figure 4:**
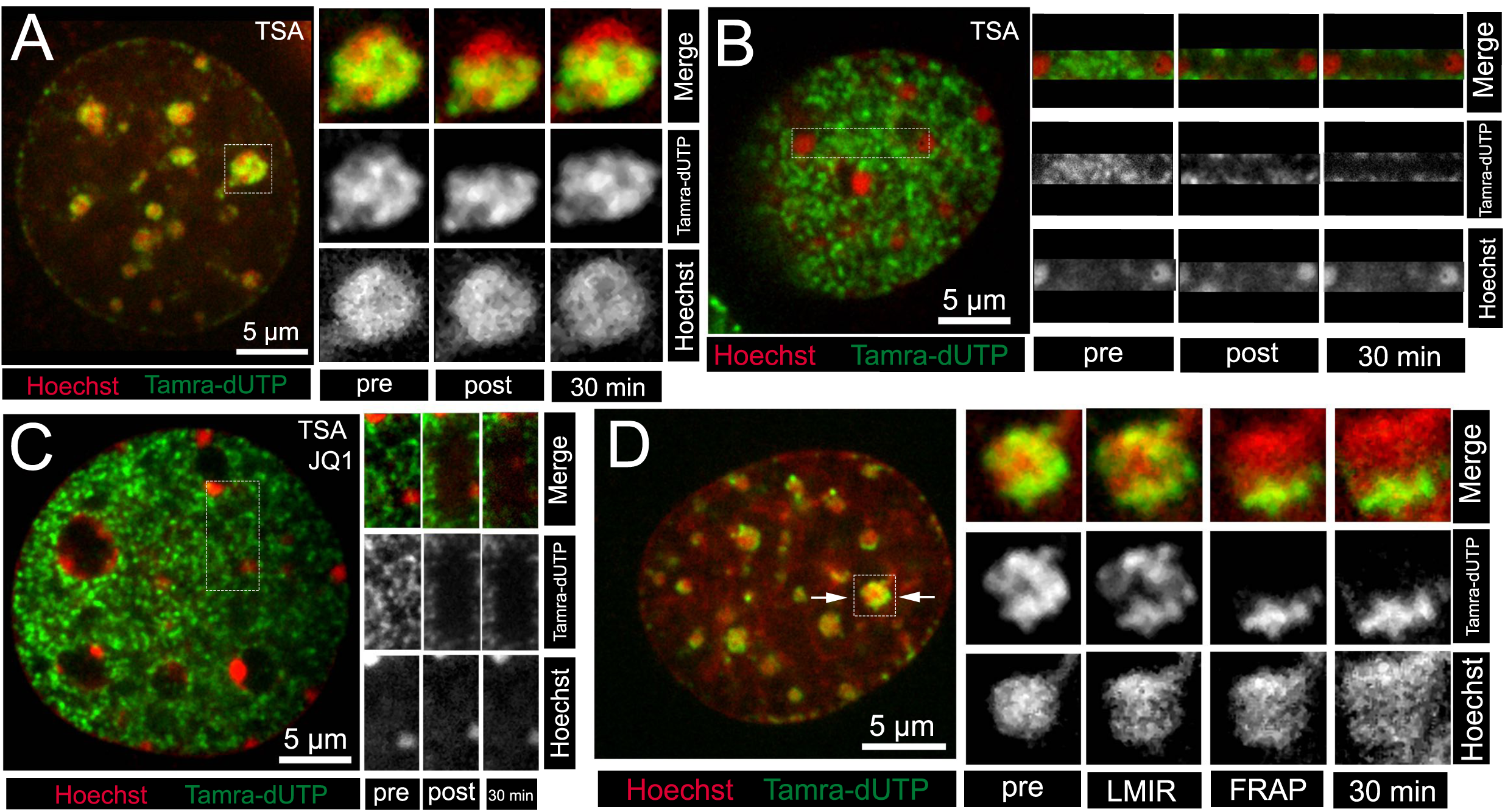
Solid-like properties of chromatin in the presence of treatments that promote dispersal of condensed intranuclear chromatin. **A:** Living C3H/10T1/2 nucleus treated with 100nM TSA for 24h stained with Hoechst (red) for DNA and showing chromatin domains labelled with TAMRA-dUTP (green). Insets of the chromocenter on the left outlined with dashes before, immediately after and 30 minutes after photobleaching. **B**: Living CH310T1/2 nucleus treated with 100nM TSA for 24h stained with Hoechst (red) for DNA and showing chromatin domains labelled with TAMRA-dUTP (green). Insets of an area full of replication foci of mid-replicating (euchromatin) from the area on the left outlined by dashes before photobleaching, immediately after FRAP and 20 minutes after photobleaching (separated by channels and merged). **C**: Photobleaching of a nucleus of a C3H/10T1/2 cell treated with 100 nM TSA and 2 mM of bromodomain inhibitor JQ-1. Replication domains still visible as distinct entities were photobleached. A magnified area full of mid-replicating replication foci (euchromatin) on the left outlined with dashes before photobleaching (separated by channels and merged), immediately FRAP and 30 minutes after photobleaching showing that the bleached replication domains are not recovering. **D**: (left) Picture of a C3H/10T1/2 cell being laser microirradiated by a 405 nm laser for laser damage induction bisecting a chromocenter (arrows). (right) Magnified chromocenter on the left outlined by dashes before laser micro-irradiation, immediately after, after photobleaching and 30 minutes after showing that the bleached replication domains are not recovering (separated by channels and merged).

One explanation for the failure of dispersed acetylated chromatin to mix is that it becomes immobilized by association with BRD-containing proteins, which have themselves been shown to undergo liquid unmixing in cells and bind to superenhancers (Benjamin et al., 2018). It was recently demonstrated that acetylated chromatin is dispersed in solution but can undergo liquid demixing in the presence of bromodomain-containing proteins (Gibson et al., 2019). This phase separation was inhibited by JQ1, an inhibitor of BRD-acetylated lysine interactions. Consequently, we tested whether treating cells with JQ1 would enable hyperacetylated chromatin to adopt liquid like behavior. We found that JQ1 had no effect on the inability of hyperacetylated chromatin to undergo mixing in FRAP experiments—the photobleached regions persisted throughout the duration of the time course (Figure 4C, Movie 13).

To further test whether dispersing chromatin can establish a liquid state, we induced rapid chromatin decondensation through laser micro-irradiation. This results in a very rapid PARP dependent decondensation of the chromatin in living cells (Strickfaden et al., 2016) and is evident in the increase in diameter and dilution of fluorescence of the Hoechst-contrasted chromatin (Figure 4D). When replication-labelled chromatin was photobleached simultaneously with laser micro-irradiation, again we did not observe mixing of the chromatin (Figure 4D, Movie 14). Thus, despite the ability of laser micro-irradiation and histone hyperacetylation to decondense and disperse native chromatin in the nucleus, they do not promote liquid-like behavior (mixing) of labelled chromatin in living cells.

### The effect of osmolarity on intranuclear chromatin organization

The expansion of chromatin following chromatin decondensation by poly(ADP-ribosyl)ation prompted us to further explore the responsiveness of chromatin structure to changes in the ion concentrations within the intracellular environment. We assessed the changes in the organization of individual chromatin fibers by transmission electron microscopy following exposure of cells to hyperosmotic and hypoosmotic conditions. Figure 5A shows that hyperosmotic treatment induced profound ultrastructural changes of the chromatin. In the corresponding binary image maps, the green indicates the positions of regions of chromatin. No longer are the ∼100 nm diameter dense chromatin structures found throughout the nucleoplasm. Instead, the chromatin accumulates in structures that are much larger than what is normally found within the nucleoplasm. Figure 5B shows a single cell nucleus that is followed through changes in osmolarity. Hyperosmolar treatment results in a marked decrease in nuclear volume while hypo-osmolar treatment results in a rapid expansion of the nuclear volume (Figure 5B). A single chromocenter is illustrated as a surface-rendered projection in Figure 5C. This reveals that, while the volume is significantly altered, the relative geometric orientation of the replication-labeled chromatin remained unchanged within individual chromocenters across these three conditions (Figure 5B). Although condensates that form by liquid-liquid phase separation usually show a sensitivity towards osmotic changes of the surrounding environment (Olins et al., 2020), the isometric contraction and expansion-behavior of the chromatin domains in this experiment resembled that of a hydrogel – not that of a liquid.

**Figure 5.**
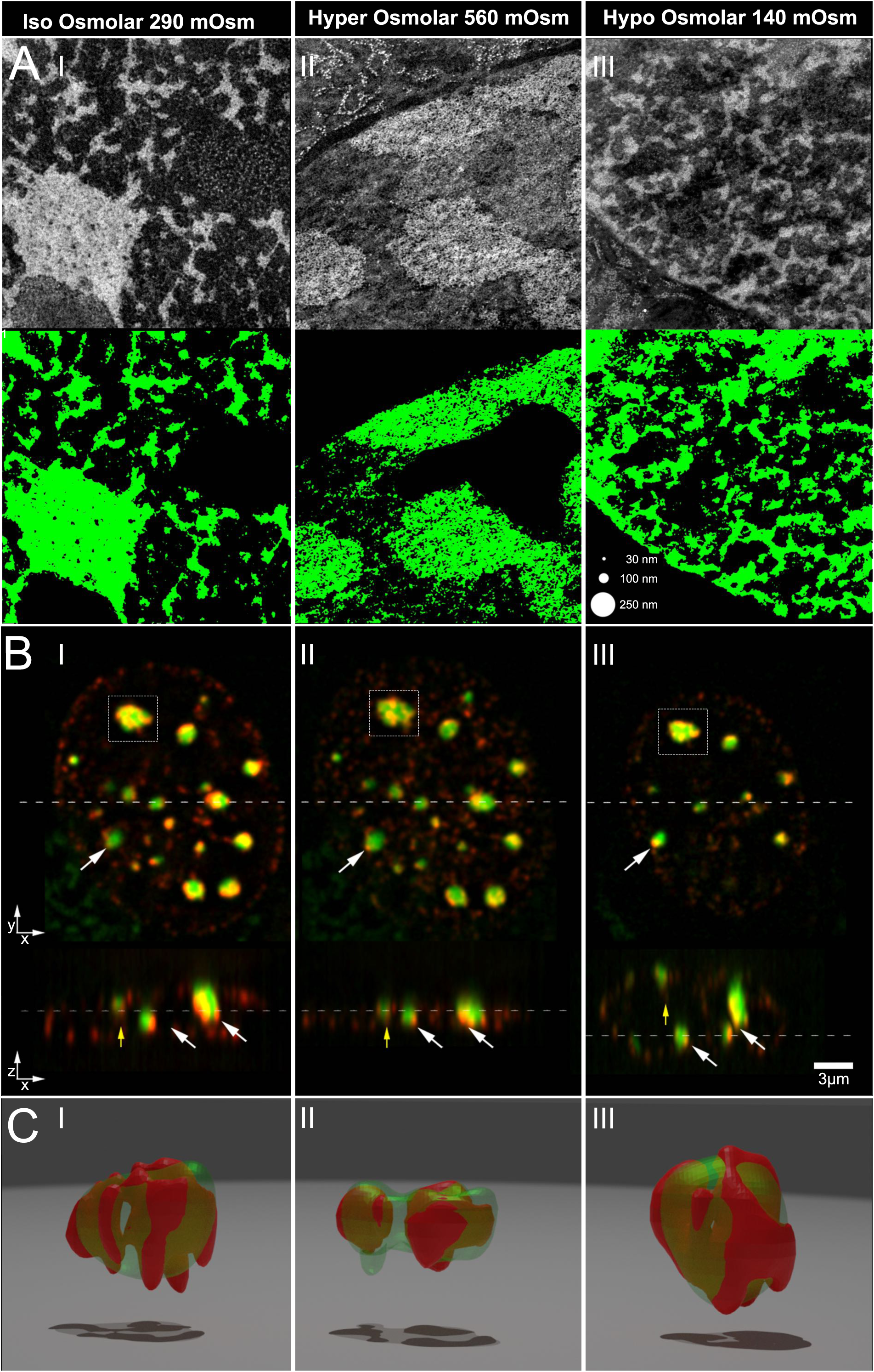
The sensitivity of condensed chromatin to changing ionic conditions in living cells. **A**: TEM micrographs showing phosphorous maps recorded by ESI of C3H10T1/2 nuclei (upper) and a binary illustration of the chromatin content (lower) in cells exposed to different osmotic environments (I iso osmolar 290 mOsm, II hyper osmolar 560 mOsm and III hypo osmolar 140 mOsm). **B:** The relative orientation of replication labeled chromatin domains remain stable irrespective of changes in the osmotic environment. The nucleus of a C3H10T1/2 nucleus replication labeled by TAMRA-dUTP (red) and KMT5C-GFP (green) exposed to different osmotic environments (I 290, II 560, III 140 mOsm) shown in a x-y (top) and a y-z (bottom) cross section. **C:** 3D reconstructions of the chromocenter outlined by dashes in the upper part of the figure.

## DISCUSSION

Interphase chromosomes exist as discrete territories that reflect short- and long-range interactions that occur primarily within the same chromatin fiber (Cremer and Cremer; Lieberman-Aiden et al., 2009). In this manner, self-interaction of the chromatin fiber across different length scales helps mold and maintain interphase chromosome structure. When assessed in vitro using short nucleosomal arrays, self-interaction leads to the formation of globular supramolecular condensates (Gibson et al., 2019; Maeshima et al., 2016b). This process is thought to recapitulate long-range chromatin interactions within chromosomes in vivo (Hansen et al., 2018; Maeshima et al., 2016b) but the physical state(s) imparted by these interactions are poorly understood. Whether chromatin condensates form a constrained solid or a mobile liquid has major implications for chromatin biology and nuclear homeostasis. Our findings, which are based on concordant in vitro and in vivo results, demonstrate that chromatin phase separates from the surrounding solution to form condensed chromatin structures with solid-like behavior. We suggest a model wherein each interphase chromosome represents a continuum of solid structures that differ in the strength of nucleosome-nucleosome interactions. The key observations that support this model are: (i) we did not observe nucleosome mobility within in vitro chromatin condensates or exchange with free chromatin in the surrounding solution; (ii) the condensates formed interacting networks when present at high concentration, but did not fuse upon contact; (iii) these assemblies were readily reversible upon removal of counterions; and (iv) in vivo imaging of DNA established an immobile and unmixed state across a range of chromatin types (e.g., heterochromatin, euchromatin) independent of histone acetylation state. Crucially, we also methodically demonstrate that recent in vitro findings of liquid behavior of chromatin (Gibson et al., 2019) result from use of nonphysiological buffer components. More physiological ionic conditions that have been vetted through decades of research on chromatin fiber self-interaction (Hansen, 2002) produce solid condensates.

Liquid-liquid phase separation has been invoked to account for multiple aspects of chromatin structure and function (Erdel and Rippe, 2018; Gibson et al., 2019; Larson et al., 2017; Liang et al., 2019; Maeshima et al., 2016a; Narlikar, 2020; Sanulli et al., 2019; Strom et al., 2017). In all cases, a core principle involves low-affinity, multivalent interactions, which often involve intrinsically disorder protein regions (IDRs). By its very nature, chromatin is a multivalent polymer that consists of intrinsically disordered histone tails that project from the surface of each nucleosome to support a range of molecular interactions. It has long been known that assembly of chromatin condensates requires these tail domains (Gordon et al., 2005; Schwarz et al., 1996), which emanate from one nucleosomal array to interact with linker DNA of other arrays (Kan et al., 2009; Kan et al., 2007). However, given the binding constant for an H4 tail peptide to DNA is 5 × 10^11^ M^-1^ (Hong et al., 1993) and that there are hundreds of thousands of nucleosomes within a condensate (Maeshima et al., 2016b), we expect the magnitude of summed interactions to confer a solid state. Importantly, each nucleosome tightly binds ∼3000 water molecules (Davey et al., 2002), leading us to propose that the extensive network of histone N-terminal tail interactions globally constrains chromatin movement to form a hydrogel.

Consistent with these properties, we found the physical state of chromatin in vivo to behave as a solid on timescale of minutes to hours, which is the relevant time scale for the functional regulation of the genome in a dividing cell. We established this by assessing the dynamic state of packaged chromatin in living cells labeled using fluorescent nucleotides and found no evidence of mixing. This applied to both the euchromatin and heterochromatin compartments, and in response to multiple stimuli that induced chromatin dispersion or decompaction where chromatin appeared to disperse isometrically to produce solid-like assemblies with decreased fiber density.

There is considerable evidence that individual gene loci undergo constrained diffusion (Chubb et al., 2002; Gasser, 2002; Marshall et al., 1997) and active transport (Chuang and Belmont, 2007; Dundr et al.; Khanna et al.). While we do not observe evidence for liquid behavior of euchromatin labelled in early S-phase, nor in cells where the chromatin has been dispersed by TSA, we do observe relative movement of the individual S-phase labelled euchromatic foci relative to each other. This relative movement has been reported before (Xiang et al., 2018). Given that each focus is thought to correspond to an individual topologically isolated domain (TAD), this would most easily be explained by flexibility of chromatin fibers that link individual TADs, as suggested previously (Xiang et al., 2018). The organization of chromatin into TADs may be responsible for the failure to observe evidence of liquidity when the chromatin fibers have been dispersed by acetylation. Thus, observing relative movement between TADs does not imply that the chromatin fibers mix, which is the defining property of liquid chromatin condensates in vitro. Given that these early S-phase euchromatic replication foci associated with TAD organization persist in cells depleted of cohesion (Cremer et al., 2019), it is also possible that lower frequency interactions between chromatin fibers mediate the solid-like behavior.

The solid state of the chromatin has at least two advantages. First, it provides mechanical stability to the genome and the cell itself, given the ability of cells to adapt to an applied force by increasing rigidity through heterochromatin formation (Stephens et al., 2019). The reduction in rigidity by histone acetylation(Maeshima et al., 2018) suggests that the viscoelastic properties of a chromatin gel can be modulated by histone post-translational modifications without transition to a liquid state. Consistent with this observation, our electron microscopy analyses revealed that the partial disassembly of condensed chromatin regions induced by histone acetylation was associated with reduced fiber-fiber interactions but did not alter the solid-like behavior. Nevertheless, a solid or gel-like state of chromatin does not preclude local nucleosome motion. Local motions of chromatin in vivo have been well documented (Maeshima et al., 2020). Small angle x-ray scattering studies suggest there is local mobility of nucleosomes within in vitro condensates that is antagonized by binding of histone H1 (Maeshima et al., 2016b). In this regard, histone H1 itself undergoes dynamic chromatin interactions in vivo and turns over every few minutes (Raghuram et al., 2009), providing a means to tune material properties, such as stiffness and elasticity, through H1 stoichiometry. Second, self-association of chromatin will compact the spatial distribution of the modifications that decorate the chromatin. Hence, by forming condensed structures, chromatin will create high local concentrations of specific histone posttranslational modifications that could seed the formation of liquid compartments. There is evidence that liquid compartments form in association with specific histone posttranslational modifications. Notably, the proteins implicated in generating these compartments, such as HP1, are not required for the formation of dense heterochromatin, as evidence by suv39h1/h2 knockout cells (Peters et al., 2001). Thus, while there is evidence for a liquid compartment associated with heterochromatin, we find that the chromatin, itself, is not in a liquid state. We therefore conclude that chromatin is a solid-like scaffold that can support the assembly of liquid-like compartments enriched in specific effector proteins that percolate throughout the fiber matrix.

## Supporting information

Supplemental Files Description and Figures

## ACKNOWLEDGEMENTS

We thank Dr. Armin Gamper for providing the JQ1 and Dr. Xuejun Sun for microscopy support. We also thank Darin McDonald and Kristal Missiaen for technical support. This work was supported by grant from the National Science Foundation (MCB-1814012) to JCH and the Canadian Institutes of Health Research to MJH (CIHR PS 162153).

## MATERIALS AND METHODS

### TAMRA-dUTPs

Fluorescently labeled nucleotides were made by incubating TAMRA-succinimidyl ester with aminoallyl-dUTP in a bicarbonate buffer as described in (Muller et al., 2007).

### Preparation of chromatin fragments from cell nuclei and chromatin solubility assay against MgCl2

Cells nuclei were isolated by lysing cells in lysis buffer (10 mM Tris-Cl, pH 7.5, 10 mM NaCl, 3 mM MgCl2, 10mm sodium butyrate, 250 mM sucrose and 0.25% V/V of NP-40). Nuclei were washed two times with same lysis buffer and were collected by centrifugation at 5000 rpm for 10 minutes. Nuclei were resuspended at 50 A260 units per ml in MNase digestion buffer (15 mM Tris–Cl pH 7.5, 15 mM NaCl, 250 mM sucrose, 2 mM CaCl2, 60 mM KCl, 15 mM β-mercaptoethanol, 0.5 mM spermidine, 0.15 mM spermine, 0.2 mM PMSF, protease and phosphatase inhibitors) and were digested with 25 U/ml of micrococcal nuclease at 37°C for 20 min. The reaction was stopped by the addition of EGTA to 10 mM and nuclei were collected by centrifugation at 5000 rpm for 10 min. Nuclei were next resuspended in 10 mM EDTA for 30 min on ice, which resulted in nuclear lysis and the release of chromatin fragments into the medium. The EDTA soluble chromatin was separated from insoluble nuclear material by centrifugation at 10000 rpm for 15 min. The isolated soluble chromatin was dialyzed overnight against 1 mM Tris-Cl (pH 8.0) and 0.1 mM EDTA at 4°C. Soluble chromatin was collected after dialysis and this was incubated in defined concentrations of MgCl2 at 4°C. Chromatin solubility was determined by leaving a solution overnight at 4°C in buffer (1 mM Tris-Cl (pH 8.0) and 0.1 mM EDTA) in the presence of MgCl2 and then centrifuge at 12500 rpm for 15 min. The optical density of the supernatant at 260 nm was used to estimate chromatin solubility.

### Formation of chromatin condensates

Soluble chromatin was stained with 1 μg/ml Hoechst and precipitated by adding 5mM MgCl2 and 100 mM KCl. The aggregation process was observed on a spinning disk microscope using the 405 nm laser at a framerate of 1 fps by placing 20 μl of the solution containing the chromatin fragments onto a glass bottom MatTek Dish and adding MgCl2 to a final concentration of 5 mM.

### Histone Octamer Purification

Recombinant Xenopus histones were purified and purchased from the CSU biochemistry protein purification facility. Purification of unlabeled histone octamers was performed by suspension of lyophilized histones H2A, H2B, H3, and H4 in unfolding buffer for >2hrs, mixing at equal molar concentrations, dialyses in 2M NaCl refolding buffer, and purification via size exclusion chromatography as previously described (Rogge et al., 2013). Fractions were run on an SDS page gel and those with equimolar amounts of the core histones were combined and concentrated to >5mg/ml.

### Alexa-488 maleimide histone H4 E63C labeling and octamer formation

Fluorophore labeling of histone H4 E63C was performed by adding an equal molar concentration of Alexa-488 maleimide to H4 E63C in unfolding buffer pH 7.0 in the presence of 0.7mM TCEP and incubated overnight in the dark at 4°C. The fluorescently tagged histone was then combined with suspended histones H2A, H2B, and H3 C110A in unfolding buffer, dialyzed in 2M NaCl and histone octamer was purified via size exclusion chromatography as described above. Histone H3 C110A was used to prevent potential labeling of H3 with the Alexa-488 fluorophore during the procedure (Gibson et al., 2019). Histones H3 E63C in combination with H3 C110A is commonly used for labeling of the histone octamer. H3 C110A has not previously been reported to affect nucleosome structure or positioning (Shimko et al., 2013).

### Assembly of Alexa-488 Labeled Nucleosomal Arrays

Nucleosomal arrays were generated using the 601-207 bp x 12 DNA template as previously described (Rogge et al., 2013). Histone octamers were combined at a ratio of 1:20 labeled to unlabeled octamer. All arrays were validated to be at or near saturation (27-28 S) using analytical ultracentrifugation as previously described (Rogge et al., 2013). Arrays were stored in a final buffer of 10mM Tris pH 7.8, 2.5 mM NaCl, 0.25 mM EDTA after dialysis.

### Chromatin condensate formation and Fluorescence Microscopy/photobleaching

Chromatin condensates were formed by first preparing the buffer solutions containing either Tris-Cl pH 7.8 or Tris-(OAc) pH 7.5 and the designated salts, 5% [w/v] glycerol, 0.1 μg/mL BSA, or 5 mM DTT. Arrays were then added to the solution to a final nucleosome concentration of 10 nM, rapidly mixed, and incubated for 20 min at 23°C. The samples where then added to MatTek glass bottom 35 mm petri dishes with a 14 mm Microwell No. 1.5 cover glass and spun at 1,000g for 3 minutes. Chromatin condensates were imaged and subject to photobleaching using an Olympus IX81 spinning disk confocal microscope, a 100x/1.40 numerical aperture objective, and a Photometrics Cascade II camera. Time lapse images were collected every 10 sec. Imaging software Slidebook6 (3I, Denver, CO) was used for image capture and analysis. Representative images in Figure 1 were obtained from zoomed in crops of the unfiltered time lapse videos.

### Cell Culture

#### Saponin labeling

In order to bulk label cells with TAMA-dUTP for chromatin fractionation (see above), cells were treated with a wash buffer consisting of 20mM HEPES pH 7.5, 138mM, KCl, 4mM MgCl2 and 3mM EGTA. After removing the wash buffer, a permeabilization buffer (wash buffer plus 0.04 μg/ml saponin) was added to the cells for 60 seconds. After carefully removing the permeabilization buffer, the wash buffer containing TAMRA-dUTP (50:1) was added carefully to the cells for 10 min. Subsequently the cells were incubated overnight in fresh DMEM.

#### Scratch labeling

Cells were grown in DMEM medium on glass bottom MatTek® dishes to a confluence of approx. 80%. The medium was completely removed from the cells and DMEM containing TAMRA-dUTP (50:1 dilution) was added so that all cells on the glass bottom were covered. The cell lawn was scratched with a 25-gauge hypodermic injection needle in parallel lines from one side to the other side. The dish was rotated 90 degrees and the cell were scratched a second time as described above. After 2 minutes the cells were washed with 1 x PBS and incubated in fresh DMEM until further usage (Schermelleh et al., 2001).

#### Inhibitor treatment

For the hyperacetylation experiments, the cells were incubated for 24h in 100 nM TSA before they were used in live-cell experiments. To inhibit the BRD-acetylated lysine interactions, we have added the inhibitor JQ1 at a concentration of 2 mM to the cells 24 h before they were used in live-cell experiments.

#### Transfection

C3H10T1/2 cells were transfected with expression plasmids the night before the experiments using Effectene in the following way: 100μl Transfection buffer, 3μl Enhancer 5 min, 4μl DNA (400ng/μl) 5 min, 6μl Effectene 20 min.

#### Fluorescence microscopy (Fractionated Chromatin/in-vivo)

20 min prior to observation, cells were stained with 0.5 μg/ml of Hoechst 33342. Photobleaching, laser micro-irradiation and time-lapse experiments were carried out on a spinning disk microscope consisting of a Zeiss Axiovert 200M inverted microscope attached to a PerkinElmer Ultraview spinning-disk confocal unit equipped with 405, 488, and 561 nm solid state laser lines, a photokinesis module, and a Photometrics PRISM BSI camera controlled by Volocity 6.3. A Zeiss 100x 1.4 NA apochromat oil immersion objective lens was used. During the experiments, cells were kept at 37°C in a humidified atmosphere containing 5% CO2. Cells were able to divide normally after photobleaching Movie 15. TAMRA-labeled chromatin was photobleached and laser microirradiated using the photokinesis module by drawing a region of interest (ROI) onto the nuclear areas, which were then exposed to localized laser illumination to photobleach or microirradiate the desired region. 66% power of the 561 nm solid state laser-line and 10 iterations were used to photobleach TAMRA-labeled chromatin; 50% power of the 488 nm solid state laser-line and 10 iterations were used to photobleach the Emerald fluorescent protein. For the laser micro-irradiation experiments, 20% power of the 405nm solid state laser and 10 iterations were used to induce DNA damage. Recovery curves (of the bleached) and depletion curves (of the non-bleached) part of the chromocenters shown in Fig S2 were created after registering the whole nucleus with the ImageJ plug-in StackReg and then measuring the mean intensity values within the photobleached (/nonbleached parts of the chromocenter) area and the background outside of the nucleus over all pictures of the time lapse experiment using the ROI-Manager of ImageJ and its function “Multi Measure”. The intensity of the measured areas of interest was then corrected by subtracting the measured background and normalized to the pre-bleaching value. The curves were plotted in Graphpad Prism. Recovery curves (of the bleached) and depletion curves (of the non-bleached) part of the chromocenters shown in Fig S2 were created after registering the whole nucleus with the ImageJ plug-in StackReg and then measuring the mean intensity values within the photobleached (/nonbleached parts of the chromocenter) area and the background outside of the nucleus over all pictures of the time lapse experiment using the ROI-Manager of ImageJ and its function “Multi Measure”. The intensity of the measured areas of interest was then corrected by subtracting the measured background and normalized to the pre-bleaching value. The curves were plotted in Graphpad Prism. Labelled cells did not show evidence of increased DNA damage (SFig. 3, Movie 16).

#### Hyper- and Hypotonic treatment of cells

Hypertonic treatment was applied to the cells by mixing the medium in the MatTek dish with a concentrated salt solution (10x PBS) 1:10 as described in (Albiez et al 2006) to reach a concentration of 560 mOsm. Hypotonic treatment was applied by adding deionized, distilled water to the medium.

#### Electron Spectroscopic Imaging

ESI was performed at a JEOL 2100F transmission electron microscope with a LaB6 filament operating at 200 kV. A Tridiem GIF post column spectrometer (Gatan) was used to record the elemental maps of phosphorus and nitrogen. Phosphorus maps were created by normalizing the background signal in void regions of the post-edge images to the background signal in pre-edge images and then subtracting the pictures. Nitrogen maps were processed by simply using the ratio method as described in (Strickfaden et al., 2015). Noise in the elemental maps was decreased by using a median filter with the radius 1 in ImageJ.

### Data-processing

#### Volocity 6.3

Data acquired at the spinning disc confocal microscope were exported as ome.tiff for further processing in FIJI or Imaris. Spinning disk confocal image stacks were deconvolved in Volocity 6.3 using a theoretical point spread function and an iterative algorithm.

#### FIJI

Motion compensation of whole nuclei was corrected using the plug-in “Stack-Reg” (Thevenaz et al., 1998). FRAP-Curves were generated using the ROI-Manager and its “Multi-Measure” function.

#### Imaris® 9.5

Imaris was used to create supporting online movies or 3D reconstructions from the recorded image stacks. The Volumes of the chromocenters and its replication labeled chromatin domains were carried out by using 3D stacks of and fitting volumes of these structures with the “Surfaces” tool. The respective surfaces were then exported in the .wrl format for further processing in Blender.

#### Blender 2.8

VRML2 files containing surfaces of chromocenters generated by Imaris® 9.5 were imported in Blender and ray-traced by the cycles render engine.

#### Digital Micrograph®

TEM pictures were recorded using Digital Micrograph®

#### Photoshop

Figures were assembled using Adobe Photoshop.

### Materials

#### Used cell-lines

C3H10T1/2 ATCC®CCL-226

#### Used expression constructs

KMT5C-Emerald (Underhill-Lab) CBX5-Emerald (Underhill-Lab)

#### Chemicals

TAMRA succinimidyl ester (5-Carboxy-tetramethylrhodamine N-succinimidyl ester) Sigma Aldrich. CAS Number 150810-68-7 PubChem Substance ID 57651396 aminoallyl-dUTP Sigma Aldrich CAS Number 936327-10-5 Trichostatin-A Sigma Aldrich CAS Number: 58880-19-6 PubChem Substance ID: 24900542 JQ1 CAS Number: 1268524-71-5 PubChem Substance ID 329825942

## References

Benjamin, R.S., Alessandra, D.A., Ann, B., Isaac, A.K., Eliot, L.C., Krishna, S., Brian, J.A., Nancy, M.H., Alicia, V.Z., John, C.M., et al. (2018). Coactivator condensation at super-enhancers links phase separation and gene control. Science 361, eaar3958.

Bian, Q., and Belmont, A.S. (2012). Revisiting higher-order and large-scale chromatin organization. Curr Opin Cell Biol 24, 359–366.

Chuang, C.H., and Belmont, A.S. (2007). Moving chromatin within the interphase nucleus-controlled transitions? Semin Cell Dev Biol 18, 698–706.

Chubb, J.R., Boyle, S., Perry, P., and Bickmore, W.A. (2002). Chromatin Motion Is Constrained by Association with Nuclear Compartments in Human Cells. Curr Biol 12, 439–445.

Cremer, M., Brandstetter, K., Maiser, A., Rao, S.S.P., Schmid, V., Mitra, N., Mamberti, S., Klein, K.-N., Gilbert, D.M., Leonhardt, H., et al. (2019). Cohesin depleted cells pass through mitosis and reconstitute a functional nuclear architecture. Biorxiv, 816611.

Cremer, T., and Cremer, C. (2001). Chromosome territories, nuclear architecture and gene regulation in mammalian cells. Nat Rev Genet 2, 292–301.

Davey, C.A., Sargent, D.F., Luger, K., Maeder, A.W., and Richmond, T.J. (2002). Solvent mediated interactions in the structure of the nucleosome core particle at 1.9 a resolution. J Mol Biol 319, 1097–1113.

Dundr, M., Ospina, J.K., Sung, M.H., John, S., Upender, M., Ried, T., Hager, G.L., and Matera, A.G. (2007). Actin-dependent intranuclear repositioning of an active gene locus in vivo. J Cell Biol 179, 1095–1103.

Eltsov, M., Maclellan, K.M., Maeshima, K., Frangakis, A.S., and Dubochet, J. (2008). Analysis of cryo-electron microscopy images does not support the existence of 30-nm chromatin fibers in mitotic chromosomes in situ. Proc Natl Acad Sci U S A 105, 19732–19737.

Erdel, F., Rademacher, A., Vlijm, R., Tunnermann, J., Frank, L., Weinmann, R., Schweigert, E., Yserentant, K., Hummert, J., Bauer, C., et al. (2020). Mouse Heterochromatin Adopts Digital Compaction States without Showing Hallmarks of HP1-Driven Liquid-Liquid Phase Separation. Mol Cell 78, 236–249 e237.

Erdel, F., and Rippe, K. (2018). Formation of Chromatin Subcompartments by Phase Separation. Biophys J 114, 2262–2270.

Ferreira, J., Paolella, G., Ramos, C., and Lamond, A.I. (1997). Spatial organization of large-scale chromatin domains in the nucleus: a magnified view of single chromosome territories. J Cell Biol 139, 1597–1610.

Finch, J.T., and Klug, A. (1976). Solenoidal model for superstructure in chromatin. Proc Natl Acad Sci U S A 73, 1897–1901.

Fussner, E., Ching, R.W., and Bazett-Jones, D.P. (2011). Living without 30nm chromatin fibers. Trends Biochem Sci 36, 1–6.

Fussner, E., Strauss, M., Djuric, U., Li, R., Ahmed, K., Hart, M., Ellis, J., and Bazett-Jones, D.P. (2012). Open and closed domains in the mouse genome are configured as 10-nm chromatin fibres. EMBO Rep 13, 992–996.

Gasser, S.M. (2002). Visualizing chromatin dynamics in interphase nuclei. Science 296, 1412–1416.

Gibson, B.A., Doolittle, L.K., Schneider, M.W.G., Jensen, L.E., Gamarra, N., Henry, L., Gerlich, D.W., Redding, S., and Rosen, M.K. (2019). Organization of Chromatin by Intrinsic and Regulated Phase Separation. Cell 179, 470–484 e421.

Gordon, F., Luger, K., and Hansen, J.C. (2005). The core histone N-terminal tail domains function independently and additively during salt-dependent oligomerization of nucleosomal arrays. J Biol Chem 280, 33701–33706.

Hansen, J.C. (2002). Conformational dynamics of the chromatin fiber in solution: determinants, mechanisms, and functions. Annu Rev Biophys Biomol Struct 31, 361–392.

Hansen, J.C., Connolly, M., McDonald, C.J., Pan, A., Pryamkova, A., Ray, K., Seidel, E., Tamura, S., Rogge, R., and Maeshima, K. (2018). The 10-nm chromatin fiber and its relationship to interphase chromosome organization. Biochem Soc Trans 46, 67–76.

Hendzel, M.J., Boisvert, F., and Bazett-Jones, D.P. (1999). Direct visualization of a protein nuclear architecture. Mol Biol Cell 10, 2051–2062.

Hong, L., Schroth, G.P., Matthews, H.R., Yau, P., and Bradbury, E.M. (1993). Studies of the DNA binding properties of histone H4 amino terminus. Thermal denaturation studies reveal that acetylation markedly reduces the binding constant of the H4 “tail” to DNA. J Biol Chem 268, 305–314.

Hyman, A.A., Weber, C.A., and Julicher, F. (2014). Liquid-liquid phase separation in biology. Annu Rev Cell Dev Biol 30, 39–58.

Joti, Y., Hikima, T., Nishino, Y., Kamada, F., Hihara, S., Takata, H., Ishikawa, T., and Maeshima, K. (2012). Chromosomes without a 30-nm chromatin fiber. Nucleus 3, 404–410.

Kan, P.Y., Caterino, T.L., and Hayes, J.J. (2009). The H4 tail domain participates in intra- and internucleosome interactions with protein and DNA during folding and oligomerization of nucleosome arrays. Mol Cell Biol 29, 538–546.

Kan, P.Y., Lu, X., Hansen, J.C., and Hayes, J.J. (2007). The H3 tail domain participates in multiple interactions during folding and self-association of nucleosome arrays. Mol Cell Biol 27, 2084–2091.

Khanna, N., Hu, Y., and Belmont, A.S. (2014). HSP70 transgene directed motion to nuclear speckles facilitates heat shock activation. Curr Biol 24, 1138–1144.

Larson, A.G., Elnatan, D., Keenen, M.M., Trnka, M.J., Johnston, J.B., Burlingame, A.L., Agard, D.A., Redding, S., and Narlikar, G.J. (2017). Liquid droplet formation by HP1alpha suggests a role for phase separation in heterochromatin. Nature 547, 236–240.

Liang, W., Yifei, G., Xiangdong, Z., Cuifang, L., Shuangshuang, D., Ru, L., Guanwei, Z., Yixuan, W., Hongyuan, Q., Yuhan, L., et al. (2019). Histone Modifications Regulate Chromatin Compartmentalization by Contributing to a Phase Separation Mechanism. Mol Cell 76, 646-659.e646.

Lieberman-Aiden, E., van Berkum, N.L., Williams, L., Imakaev, M., Ragoczy, T., Telling, A., Amit, I., Lajoie, B.R., Sabo, P.J., Dorschner, M.O., et al. (2009). Comprehensive mapping of long-range interactions reveals folding principles of the human genome. Science 326, 289–293.

Maeshima, K., Hihara, S., and Takata, H. (2010). New insight into the mitotic chromosome structure: irregular folding of nucleosome fibers without 30-nm chromatin structure. Cold Spring Harb Symp Quant Biol 75, 439–444.

Maeshima, K., Ide, S., Hibino, K., and Sasai, M. (2016a). Liquid-like behavior of chromatin. Curr Opin Genet Dev 37, 36–45.

Maeshima, K., Rogge, R., Tamura, S., Joti, Y., Hikima, T., Szerlong, H., Krause, C., Herman, J., Seidel, E., DeLuca, J., et al. (2016b). Nucleosomal arrays self-assemble into supramolecular globular structures lacking 30-nm fibers. The EMBO Journal 35.

Maeshima, K., Tamura, S., Hansen, J.C., and Itoh, Y. (2020). Fluid-like chromatin: Toward understanding the real chromatin organization present in the cell. Curr Opin Cell Biol 64, 77–89.

Maeshima, K., Tamura, S., and Shimamoto, Y. (2018). Chromatin as a nuclear spring. Biophysics Physicobiology 15, 189–195.

Marshall, W.F., Straight, A., Marko, J.F., Swedlow, J., Dernburg, A., Belmont, A., Murray, A.W., Agard, D.A., and Sedat, J.W. (1997). Interphase chromosomes undergo constrained diffusional motion in living cells. Curr Biol 7, 930–939.

Nakashima, K.K., Vibhute, M.A., and Spruijt, E. (2019). Biomolecular Chemistry in Liquid Phase Separated Compartments. Front Mol Biosci 6, 21.

Narlikar, G.J. (2020). Phase-separation in chromatin organization. J Biosci 45.

Olins, A.L., Gould, T.J., Boyd, L., Sarg, B., and Olins, D.E. (2020). Hyperosmotic stress: in situ chromatin phase separation. Nucleus 11, 1–18.

Otterstrom, J., Castells-Garcia, A., Vicario, C., Gomez-Garcia, P.A., Cosma, M.P., and Lakadamyali, M. (2019). Super-resolution microscopy reveals how histone tail acetylation affects DNA compaction within nucleosomes in vivo. Nucleic Acids Res 47, 8470–8484.

Ou, H.D., Phan, S., Deerinck, T.J., Thor, A., Ellisman, M.H., and O’Shea, C.C. (2017). ChromEMT: Visualizing 3D chromatin structure and compaction in interphase and mitotic cells. Science 357.

Perry, M., and Chalkley, R. (1982). Histone acetylation increases the solubility of chromatin and occurs sequentially over most of the chromatin. A novel model for the biological role of histone acetylation. J Biol Chem 257, 7336–7347.

Peters, A.H., O’Carroll, D., Scherthan, H., Mechtler, K., Sauer, S., Schofer, C., Weipoltshammer, K., Pagani, M., Lachner, M., Kohlmaier, A., et al. (2001). Loss of the Suv39h histone methyltransferases impairs mammalian heterochromatin and genome stability. Cell 107, 323–337.

Ricci, Maria A., Manzo, C., García-Parajo, M.F., Lakadamyali, M., and Cosma, Maria P. (2015). Chromatin Fibers Are Formed by Heterogeneous Groups of Nucleosomes In Vivo. Cell 160, 1145–1158.

Rogge, R.A., Kalashnikova, A.A., Muthurajan, U.M., Porter-Goff, M.E., Luger, K., and Hansen, J.C. (2013). Assembly of nucleosomal arrays from recombinant core histones and nucleosome positioning DNA. J Vis Exp.

Sanulli, S., Trnka, M.J., Dharmarajan, V., Tibble, R.W., Pascal, B.D., Burlingame, A.L., Griffin, P.R., Gross, J.D., and Narlikar, G.J. (2019). HP1 reshapes nucleosome core to promote phase separation of heterochromatin. Nature 575, 390–394.

Schermelleh, L., Solovei, I., Zink, D., and Cremer, T. (2001). Two-color fluorescence labeling of early and mid-to-late replicating chromatin in living cells. Chromosome Res 9, 77–80.

Schwarz, P.M., Felthauser, A., Fletcher, T.M., and Hansen, J.C. (1996). Reversible oligonucleosome self-association: dependence on divalent cations and core histone tail domains. Biochemistry 35, 4009–4015.

Shimko, J.C., Howard, C.J., Poirier, M.G., and Ottesen, J.J. (2013). Preparing semisynthetic and fully synthetic histones h3 and h4 to modify the nucleosome core. Methods Mol Biol 981, 177–192.

Stephens, A.D., Liu, P.Z., Kandula, V., Chen, H., Almassalha, L.M., Herman, C., Backman, V., O’Halloran, T., Adam, S.A., Goldman, R.D., et al. (2019). Physicochemical mechanotransduction alters nuclear shape and mechanics via heterochromatin formation. Mol Biol Cell 30, 2320–2330.

Strick, R., Strissel, P.L., Gavrilov, K., and Levi-Setti, R. (2001). Cation-chromatin binding as shown by ion microscopy is essential for the structural integrity of chromosomes. J Cell Biol 155, 899–910.

Strickfaden, H., McDonald, D., Kruhlak, M.J., Haince, J.F., Th’ng, J.P., Rouleau, M., Ishibashi, T., Corry, G.N., Ausio, J., Underhill, D.A., et al. (2016). Poly(ADP-ribosyl)ation-dependent Transient Chromatin Decondensation and Histone Displacement following Laser Microirradiation. J Biol Chem 291, 1789–1802.

Strickfaden, H., Missiaen, K., Hendzel, M.J., and Underhill, D.A. (2019). KMT5C displays robust retention and liquid-like behavior in phase separated heterochromatin. Biorxiv, 776625.

Strickfaden, H., Xu, Z.Z., and Hendzel, M.J. (2015). Visualization of miniSOG Tagged DNA Repair Proteins in Combination with Electron Spectroscopic Imaging (ESI). J Vis Exp.

Strom, A.R., Emelyanov, A.V., Mir, M., Fyodorov, D.V., Darzacq, X., and Karpen, G.H. (2017). Phase separation drives heterochromatin domain formation. Nature 547, 241–245.

Thevenaz, P., Ruttimann, U.E., and Unser, M. (1998). A pyramid approach to subpixel registration based on intensity. IEEE Trans Image Process 7, 27–41.

Thoma, F., Koller, T., and Klug, A. (1979). Involvement of histone H1 in the organization of the nucleosome and of the salt-dependent superstructures of chromatin. J Cell Biol 83, 403–427.

Xiang, W., Roberti, M.J., Heriche, J.K., Huet, S., Alexander, S., and Ellenberg, J. (2018). Correlative live and super-resolution imaging reveals the dynamic structure of replication domains. J Cell Biol 217, 1973–1984.

Young Lee, J., and Hirose, M. (2014). Effects of Disulfide Reduction on the Emulsifying Properties of Bovine Serum Albumin. Biosci Biotechnology Biochem 56, 1810–1813.

