## Supplemental Files Description and Figures for "Condensed chromatin behaves like a solid on the mesoscale in vitro and in living cells"

### Supporting online material

#### Supporting Online Figures:

SFig1 **A:** Fluorescence recovery after photobleaching of entire chromatin condensates formed by Alexa488 labeled nucleosomal arrays in 4mM MgCl2, BSA, and DTT (Scale bar, 1µm). **B:** Graphical representation of percent fluorescence recovery after photobleaching of condensates from Figure 1 c-f (n=4 condensates per condition).

SFig2: **Photobleaching of KMT5C-EM and CBX5-EM after photobleaching significant parts of a chromocenter.**

**A:** Photobleaching of a region of a chromocenter in KMT5C-EM transfected cells shows slow recovery of the protein in the bleached area (red line). Symmetric loss of fluorescence in the non-bleached part of the chromocenter shows decline in signal caused by liquid-like distribution within the chromocenter into the photobleached areas (green line). **B:** Photobleaching of parts in a chromocenter in CBX-5-EM transfected cells shows rapid recovery in the bleached area (red) and an initial loss of signal in the unbleached portion of chromocenter showing less pronounced liquid properties.

SFig3**: Testing for DNA damage by replication labeling.**

Plotting average intensities of gH2AX and dUTP-TAMRA signals within nuclei shows very low correlations between DNA-damage and replication label intensity for cells labeled **A:** 12h and **B:** 24h respectively.

#### Supporting online Movies:

Movie 01: Precipitation of fractionated chromatin after adjusting the buffer to 5mM $MgCl_2$. This movie shows timelapse results obtained for figure 1B.

Movie 02: Photobleaching of reconstituted fluorescent chromatin arrays in TrisAc MgOAc KOAc with Glycerol DTT and BSA. This movie shows timelapse results obtained for figure 1I.

Movie 03: Photobleaching of reconstituted fluorescent chromatin arrays in TrisAc MgOAc KOAc only. This movie shows timelapse results obtained for figure 1J.

Movie 04: Photobleaching of reconstituted fluorescent chromatin arrays in TrisAc MgOAc KOAc with DTT and BSA only. This movie shows timelapse results obtained for figure 1K.

Movie 05: Photobleaching of reconstituted fluorescent chromatin arrays in TrisAc MgOAc KOAc with BSA only. This movie shows timelapse results obtained for figure 1L.

Movie 06: Photobleaching of reconstituted fluorescent chromatin arrays in TrisAc MgOAc KOAc with Glycerol and DTT only. This movie shows timelapse results obtained for figure 1M.

Movie 07: Photobleaching of replication labeled fluorescent chromatin arrays in a chromocenter. This movie shows timelapse results obtained for figure 2A.

Movie 08: Photobleaching of chromocenters in cells expressing KMT5C-Emerald Tamra-dUTP. This movie shows timelapse results obtained for figure 2B.

Movie 09: Photobleaching of chromocenters in cells expressing CBX5-Emerald Tamra-dUTP. This movie shows timelapse results obtained for figure 2C.

Movie 10: Euchromatin Frap of Tamra dUTP labelled replication foci. This movie shows timelapse results obtained for figure 2C.

Movie 11: Chromocenter FRAP of TSA treated cells. This movie shows timelapse results obtained for figure 4A.

Movie 12: Euchromatin FRAP of TSA treated cells. This movie shows timelapse results obtained for figure 4B.

Movie 13: Chromocenter/Euchromatin FRAP of TSA and JQ treated cells. This movie shows timelapse results obtained for figure 4C.

Movie 14: Laser micro-irradiation of chromatin followed by FRAP. This movie shows timelapse results obtained for figure 4D.

Movie 15: Survival and proliferation of C3H10T1/2 cells after chromatin FRAP in reference to materials and methods.

SFig. 1

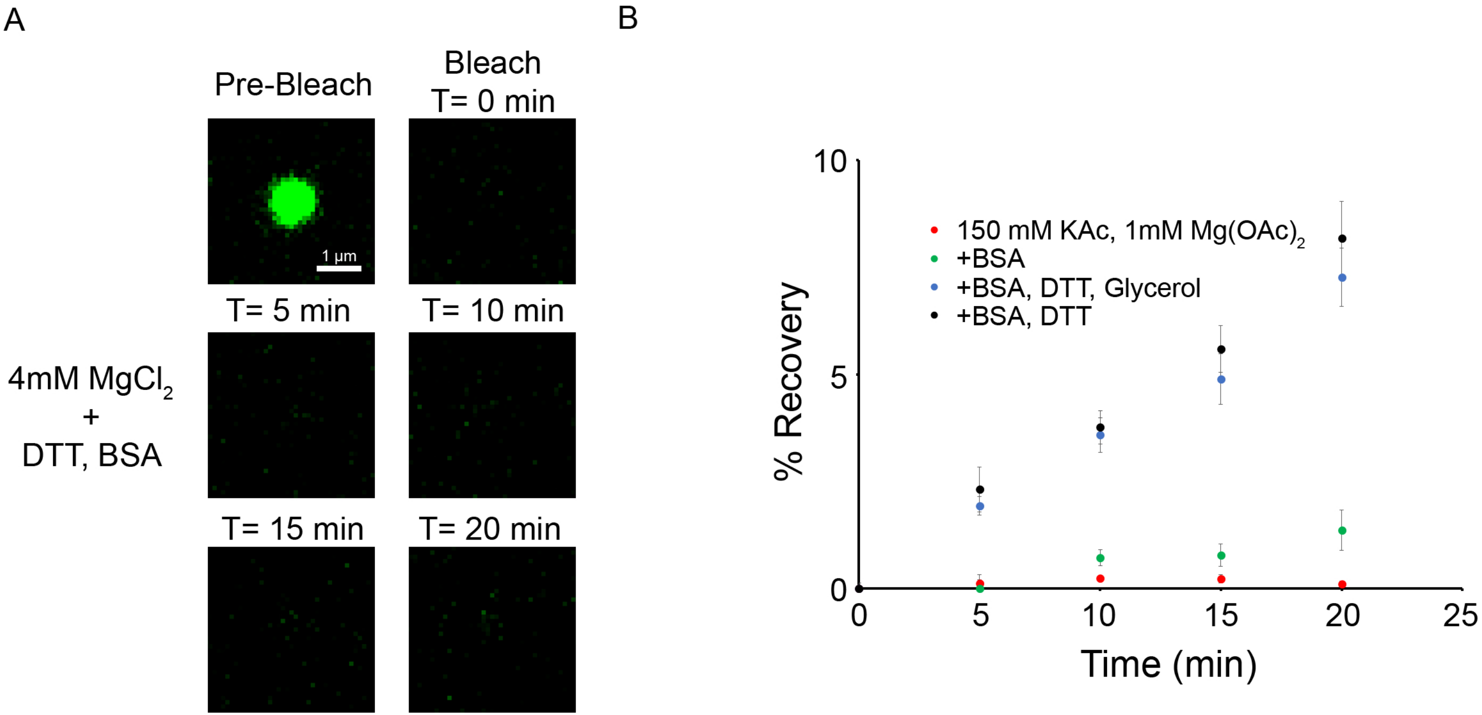
A
Pre-Bleach
Bleach
T= 0 min
1 µm
T= 5 min
T= 10 min
4mM MgCl2
+
DTT, BSA
T= 15 min
T= 20 min
B
% Recovery
10
5
0
0
5
10
15
20
25
Time (min)
150 mM KAc, 1mM Mg(OAc)2
+BSA
+BSA, DTT, Glycerol
+BSA, DTT

SFig. 2

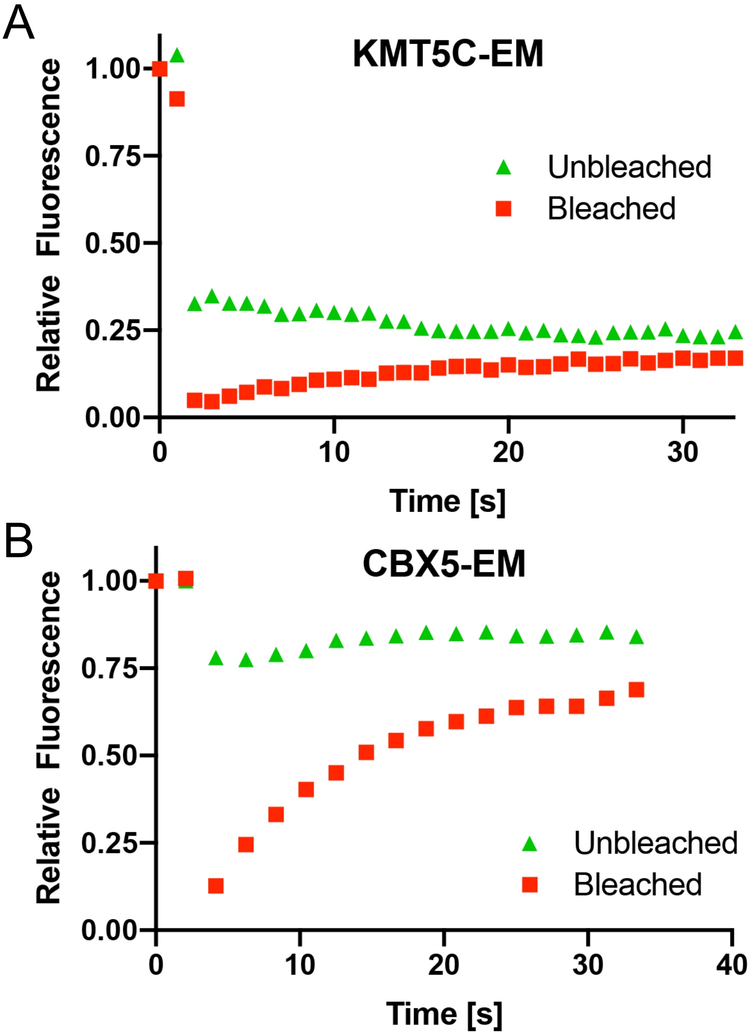
A
KMT5C-EM
Relative Fluorescence
1.00
0.75
0.50
0.25
0.00
0
10
20
30
Time [s]
Unbleached
Bleached
B
CBX5-EM
Relative Fluorescence
1.00
0.75
0.50
0.25
0.00
0
10
20
30
40
Time [s]
Unbleached
Bleached

SFig. 3

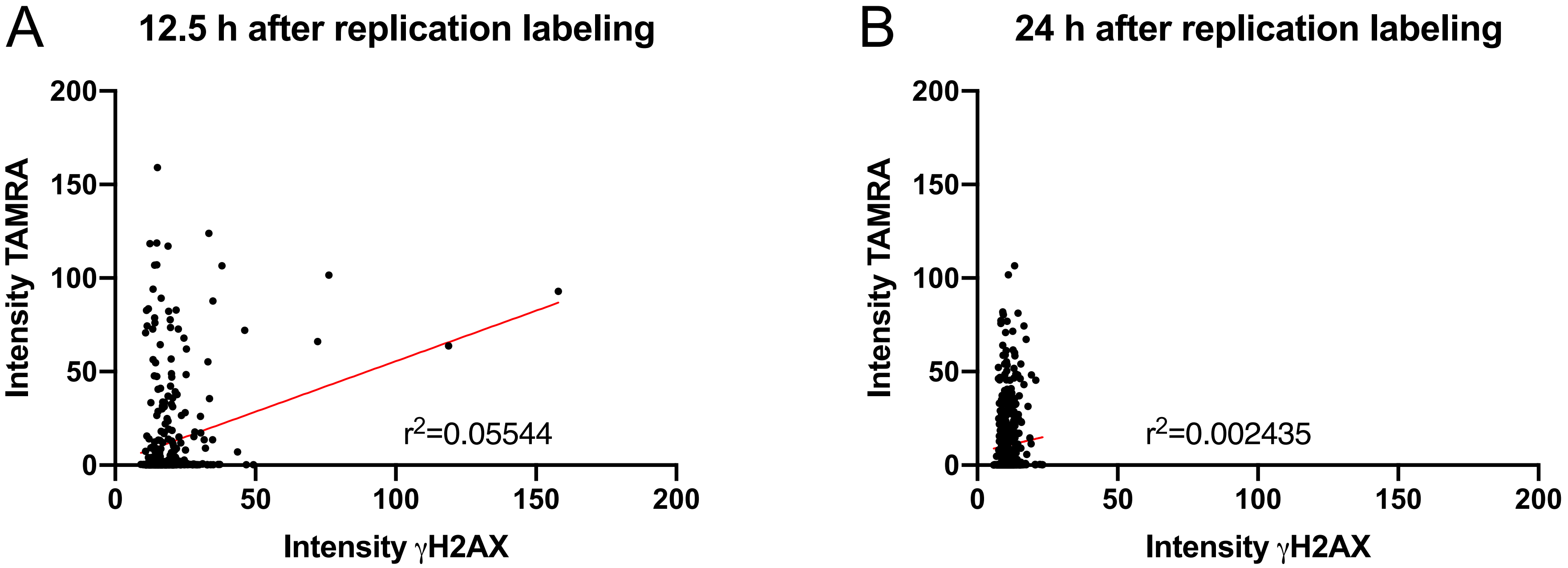

A
12.5 h after replication labeling
Intensity TAMRA
Intensity γH2AX
r2=0.05544
0
50
100
150
200
B
24 h after replication labeling
Intensity TAMRA
Intensity γH2AX
r2=0.002435
